# Suitable use of FRET-based Biosensors for Quantitative Detection of GPCR Activation

**DOI:** 10.1101/2022.06.01.494354

**Authors:** Nikolaj K. Brinkenfeldt, André Dias, Jesper M. Mathiesen, Karen L. Martinez

**Affiliations:** University of Copenhagen

**Keywords:** Biosensor, G protein-coupled receptor, cAMP, FRET, Epac, Imaging, Screening

## Abstract

Cyclic adenosine 3’,5’-monophosphate is an important second messenger molecule that regulates many downstream signaling pathways in cells. Detection of cAMP levels relies on screenings of cell lysates or the use of genetically encoded biosensors for detection in living cells. Genetically encoded biosensors are, however, primarily used for bioimaging and rarely in high-throughput screenings of potential drug candidates. Here, we describe a quantitative fluorescence-based imaging method based on measurements of single living cells. We used a genetically encoded Epac149 biosensor to investigate cAMP production in living cells following ligand stimulation. The study revealed a dependence of the measured cAMP levels on the expression level of the biosensor in transiently transfected cells. While the biosensor maintained linearity of the signal at high expression levels, the linearity of the biosensor was lost at lower expression levels due to a deficit of the biosensor compared to the maximum possible production of cAMP in the cells. This problem was circumvented by establishment of a stable cell line with constitutive expression of the biosensor. We established dose response curves by stimulation with the β_1_-adrenergic receptor partial agonist denopamine and observed up to 1.48-fold difference in the cellular response as well as up to 4.27-fold difference in LogEC_50_ values between cells with insufficient and sufficient biosensor expression. Careful characterization and control of the biosensor expression is therefore important in order to conduct quantitative analysis of the cAMP production and it allows the use of genetically encoded biosensor to be applied in high-throughput screenings.

---

Detection of cell responses upon stimulation is a necessary step to understand cell functions, and specific detection of a given cellular response provides information about the specific signaling pathway involved upon stimulation. In general, there are two ways to measure a given cellular response. The first is by detection of intracellular signaling molecules in cell lysate. Such an approach offers poor temporal resolution and no spatial resolution of the signaling molecules (Evellin, S. et al. 2004). The second approach is to use fluorescence based biosensors for detection of signaling events, which allows spatiotemporal detection in living cells (Ghigo, A. & Mika, D. 2019).

The first fluorescence-based biosensor not detecting ions such as Ca^2+^, H^+^, Na^+^, Mg^2+^, and Cl^-^, FlCRhR, was used for detection of intracellular cyclic adenosine 3’,5’-monophosphate (cAMP) in living cells and was based on detection of Protein Kinase A (PKA) dissociation when cAMP binds (Adams, S. R. et al. 1991). Being the first however, it required chemical labeling and microinjection in cells, which limited its applications. Numerous genetically encoded fluorescence based biosensors suitable for intracellular detection of signaling molecules in living cells have been developed since then, including sensors for Ca^2+^, (Miyawaki, A.. et al. 1997), various ions (Bischof, H. et al. 2017 and Oh, J. et al. 2019), detection of intracellular signaling events (Lin, W. et al. 2019), and surveillance of cell death (Bardet, P-L. et al. 2008).

There are two major types of genetically encoded biosensors. Biosensors that rely on a ligand binding domain inserted in the vicinity of the chromophore of a single fluorescent protein (Baird, G. S. et al. 1999 and Nakai, J. et al. 2001 and Nasu, Y. et al. 2021 and Mita, M. et al. 2022), and biosensors that rely on two fluorescent proteins and the energy transfer from one to another for detection (San Martin, A. et al. 2022). While single FP biosensors are small and suitable for multi-color imaging, they have a small dynamic range, low fluorescence intensity, and a sensitivity to changes in pH. The other major type of genetically encoded biosensors relies on resonance energy transfer. These biosensors typically have large dynamic ranges, good sensitivity and are robust towards changes in pH. Bioluminescence resonance energy transfer (BRET)-based biosensors do not need excitation light (Barak, L. S. et al 2008), however, the luminescence originating from BRET-sensors is typically weak when expressed in living species (Kim, N. et al. 2021). Förster resonance energy transfer (FRET)-based biosensors are less invasive but require excitation light. However, the fluorescence signals are greater compared to the luminescence of BRET-based sensors and the dynamic range of FRET-based sensors are large (Sprenger, J. U. and Nikolaev, V. O. 2013), which makes them desirable for studies in living cells. Conventionally, the FRET-pair consists of eCFP and eYFP and is expressed on each side of a ligand binding domain (Ponsioen, B. et al. 2004), but many sensors with improvements to the FRET-pair are developed.

The list of FRET-based sensors is substantially longer compared to other biosensors (Okumoto, S. et al. 2012) and includes, among many others, sensors for imaging of ions (Mank, M. et al. 2008 and Vinkenborg, J. L. et al. 2009), imaging of kinase activity (Crosby, K. C. et al. 2011 and Zhang, J. et al. 2001), and imaging of metabolites such as glucose (Takanaga, H. et al. 2008), cGMP (Niino, Y. et al. 2010), and cAMP (Nagai, T. et al. 2001 and Klarenbeek, J. B. et al. 2011).

Many G protein-coupled receptors tightly regulate the activity of adenylyl cyclase (AC) activity, which is an enzyme that catalyzes synthesis of cAMP (Serezani, C. H. et al. 2008), an important intracellular analyte involved in the control of numerous cellular functions and many pathophysiological processes in different organs (Beavo, J. A. & Brunton, L. L. 2002). GPCRs constitute the largest family of integral membrane proteins in eukaryotes. They are triggered by various types of endogenous ligands, such as peptides, neurotransmitters, and hormones, as well as exogenous ligands, such as odorants, tastants, and light (Martins et al. 2012). Today, more than 800 different structures are known to be encoded in the human genome, each with essential roles in the vast variety of biological processes that take place (Fredriksson et al. 2003). Due to their abundance and key role in many physiological processes, GPCRs are attractive pharmacological targets, and more than one-third of all FDA approved drugs target GPCRs (Hauser et al. 2017). The primary signaling transduction of GPCRs happens through heterotrimeric G proteins composed of alpha, beta, and gamma subunits (Gα, Gβ, and Gγ) (Duc, N. M et al. 2015) and exposure of GPCRs to stimuli leads to a dissociation of the G protein into an α-subunit and a β/γ-complex (Weis, W. I. and Kobilka, B. K. 2018). There are 18 distinct members of the α-subunit family (Glukhova, A. et al. 2018), from which 11 regulate (Syrovatkina, V. et al. 2016) AC activity. Therefore, cAMP is one of the major players in signal transduction, which makes cAMP assays highly relevant and abundantly used for discovery of potential drug candidates.

Conventional cAMP assays are based on detection of cellular cAMP in cell lysates by competition with a labeled cAMP for a limited number of anti-cAMP antibody binding sites (Gabriel, D. et al. 2003), and various technologies have been developed for measurements in high-throughput screenings (Williams, C. 2004 & Hill. S. J. et al. 2010). While these assays are all based on antibody detection, the protocols differ substantially (Zhang, R. and Xie, X. 2012). While some assays use enzyme complementation by release of small peptide fragments from anti-cAMP antibodies (Eglen, R. M. and Singh, R. 2003 & Weber, M. et al. 2004) other assays utilize sensitive proximity chemiluminescence (Eglen, R. M. 2008) or HTRF-based measurements to detect cAMP (Degorce, F. et al. 2009). Nonetheless, because these assays rely on antibody detection in cell lysates, they offer very poor temporal resolution and no spatial resolution (Evellin, S. et al. 2004), which limits the biological information that can be obtained. To circumvent these challenges, the use of genetically encoded biosensors has increased tremendously and today there are more than 35 distinct genetically encoded biosensors (Jiang, J. Y. et al. 2017), including sensors based on single FPs (Kitaguchi, T. et al. 2013 & Odaka, H. et al. 2014 & Harada, K. et al. 2017, BRET (Rangaraju, V. et al. 2014 and San Martin, A. et al. 2022), or FRET (Nikolaev, V. O. et al. 2004 and Klarenbeek, J. et al. 2015). Genetically encoded biosensors are mostly used for imaging, for example to investigate cAMP compartmentalization in living cells (Anton, S. E. et al. 2022 & Berisha, F. et al. 2021 & Surdo, N. C. et al. 2017), but are rarely in high-throughput screening. Only few examples are reported (Vedel, L. et al. 2015 and Tewson, P. H. et al. 2015).

Here we describe a new approach to detect activation of G_s_-and G_i/o_-coupled receptors by quantitative fluorescence-based single-cell microscopy. We used a previously described (Violin, J. D. et al. 2008) Epac1-based biosensor with another, more efficient (Rizzon, M. A. et al. 2004), FRET pair, mCerulean (mCer) and mCitrine (mCit) (Mathiesen, J. M. et al. 2013). The biosensor, named Epac149, has a large dynamic range and good sensitivity, comparable with other FRET-based sensors (Nikolaev, V. O. et al. 2004 & Norris, R. P. et al. 2009). We show that changes in intracellular cAMP levels after stimulation of the cells depends on the biosensor expression level in each cell. Typical characterization of biosensor performance is conducted in solution with an excess of the biosensor compared to cAMP. However, when confining the biosensor inside cells, the linearity of the signal is lost if the biosensor is not expressed in a quantity larger than the possible cAMP production of the cell. By accounting for the variation in biosensor expression level of each single cell, we were able to establish a minimum threshold needed to quantitatively interpret changes in intracellular cAMP levels. Additionally, we show that to adapt the use of genetically encoded biosensors to high-throughput settings, it is important to establish cell lines that express levels of biosensor above the threshold.

## Materials and methods

### Materials

Dulbecco’s modified Eagle’s medium/F-12 GlutaMAX™ (DMEM), blasticidin S HCl and hygromycin B were purchased from Fisher Scientific. TurboFect™ transfection reagent, zeocin, penicillin/streptomycin and tetracycline hydrochloride were purchased from Thermo Fisher Scientific. Fetal bovine serum (FBS) and trypsin-EDTA 0.25 % EDTA 0.02 % were purchased from InVitro. Forskolin and R(-)-Denopamine were purchased from Sigma Aldrich and SNAP-Surface AF647 was purchased from New England Biolabs. PBS buffer pH=7.4 (137 mM NaCl, 2.7 mM KCl, 10 mM Na_2_HPO_4_, 1.8 mM KH_2_PO_4_) and Epac-buffer (140 mM NaCl, 5 mM KCl, 1 mM MgCl_2_, 1 mM CaCl_2_, 10 mM HEPES, 10 mM Glucose) were homemade.

### Cell Lines

HEK293 cell line (ATTC). Flp-In™293 cells stably expressing TetOn inducible SNAP-β_1_-adrenergic receptors (iSNAP-β_1_AR) created within the group. SNAP-β_1_AR plasmid (Cisbio). Stable HEK293-Epac149 cell line created by Jesper M. Mathiesen. Epac149 biosensor plasmid produced by Jesper M. Mathiesen.

### Cell culture

HEK293, iSNAP-β_1_AR and HEK293-Epac149 cells stably expressing Epac149 biosensor were all cultured in DMEM supplemented with 10 % FBS at 37°C and 5 % CO_2_. For maintenance of stable expression in iSNAP-β_1_AR the medium was supplemented with 15 µg/mL blasticidin S HCl and 100 µg/mL hygromycin B. In the case of HEK293-Epac149, 50 µg/mL zeocin and 100 µg/mL penicillin/streptomycin was added to the culture medium.

### Expression of Epac149 biosensor in HEK293 and iSNAP-β1AR cell line or expression of SNAP-β1AR in HEK293-Epac149 cell line

Adherent cells grown to 70-90 % confluency in T25 flasks were detached by removal of growth medium and washing with PBS buffer once, followed by incubation with 2 mL trypsin-EDTA for 2 minutes. After detachment, 2 mL growth medium was added to the cells and they were then pelleted at 1200 rpm for 2 minutes. The supernatant was carefully removed, and the cells were resuspended in 2 mL growth medium. The number of cells was determined by counting in a hemocytometer. Approximately 200000 cells/well were transferred to round sterilized glass cover slips deposited in a 6-well culture plate. Each well was filled with 2 mL growth medium containing appropriate antibiotics, and incubated overnight at 37 °C and 5 % CO_2_. The following day, cells were transiently transfected with either 1 µg/mL Epac149 biosensor plasmid (HEK293 and iSNAP-β1AR cells) or 1 µg/mL SNAP-β1AR plasmid (HEK293-Epac149 cells) following the manufacturer protocol for the commercially available transfection reagent TurboFect™. The cells were cultured for 18-24 hours at 37°C and 5 % CO_2_ to allow gene expression.

### Expression of inducible SNAP-β1AR in iSNAP-β1AR cell line

Inducible SNAP-β1AR was induced simultaneously with transfection by addition of 1 µg/mL tetracycline to each sample.

### Labeling of SNAP-β1AR

Cells expressing SNAP-β1AR were labeled with 2.5 µM SNAP-Surface AF647 in DMEM by covalent SNAP tag coupling for 30 minutes at 37°C and 5 % CO_2_. The samples were washed three times with DMEM followed by two times with Epac-buffer before imaging.

### Quantitative fluorescence microscopy

Laser scanning confocal microscopy (LSCM) was performed on an Olympus IX81 confocal setup using a UPLSAPO 100x oil immersion objective with a 1.4 numerical aperture. The mCer fused to the Epac149 biosensor was excited with a wavelength of 405 nm. The emission of mCer was measured at 465 - 495 nm and the emission of mCit was measured at 535 - 555 nm. SNAP-β_1_AR was excited with a wavelength of 635 nm and emission was measured at 655 - 755 nm.

### Data analysis

All measured fluorescence intensities were extracted from raw microscopy images by custom-written MATLAB (The Mathworks, Inc., Natick, MA) scripts. Individual cells were segmented using a watershed algorithm using the SNAP-β_1_AR channel, followed by connected-component labeling. The FRET ratio was then calculated as the ratio between mCitrine and mCerulean intensities for each cell. The change in intracellular cAMP levels was calculated as the percentage decrease in the FRET ratio. Data were presented using GraphPad Prism (GraphPad Software, San Diego, CA).

## Results and discussion

### Detection of intracellular cAMP levels by quantitative fluorescence microscopy

To detect changes of cAMP in living cells while imaging, we used the Epac149 biosensor (Mathiesen, J. M. et al. 2013). In the absence of cAMP, mCer and mCit are in close proximity which allows FRET from mCer to mCit. The fluorescence intensity emitted by the sensor changes due to an increased distance between mCer and mCit upon binding of cAMP to the sensor, shown in **Figure 1B**. The fluorescence of mCit when directly excited indicated the amount of biosensor expressed in HEK293 cells transiently transfected with the biosensor. The FRET-ratio between mCer and mCit measured after direct excitation of mCer is used as a signal baseline and indicates the level of cAMP in non-stimulated cells. The FRET signal when the cells reached their maximum cAMP production was measured with forskolin, which binds directly to the AC, activates it and leads to raising of cAMP levels in cells. Here, we added 100 µM forskolin to the cells to induce maximum production of cAMP (Seamon, K. B. et al. 1981). The cells were imaged before and 15 minutes after the addition of forskolin. As shown in **Figure 1C**, in the case of twenty representative cells, we observed up to a 16-fold difference in the maximum change of intracellular cAMP levels upon stimulation with forskolin. While the intracellular cAMP levels in several cells increased to a similar value, the levels of cAMP barely increased in some other cells, which suggests an uneven response throughout the entire cell population. Such an uneven response allows only a semi-quantitative interpretation of the cellular response.

**Figure 1:**
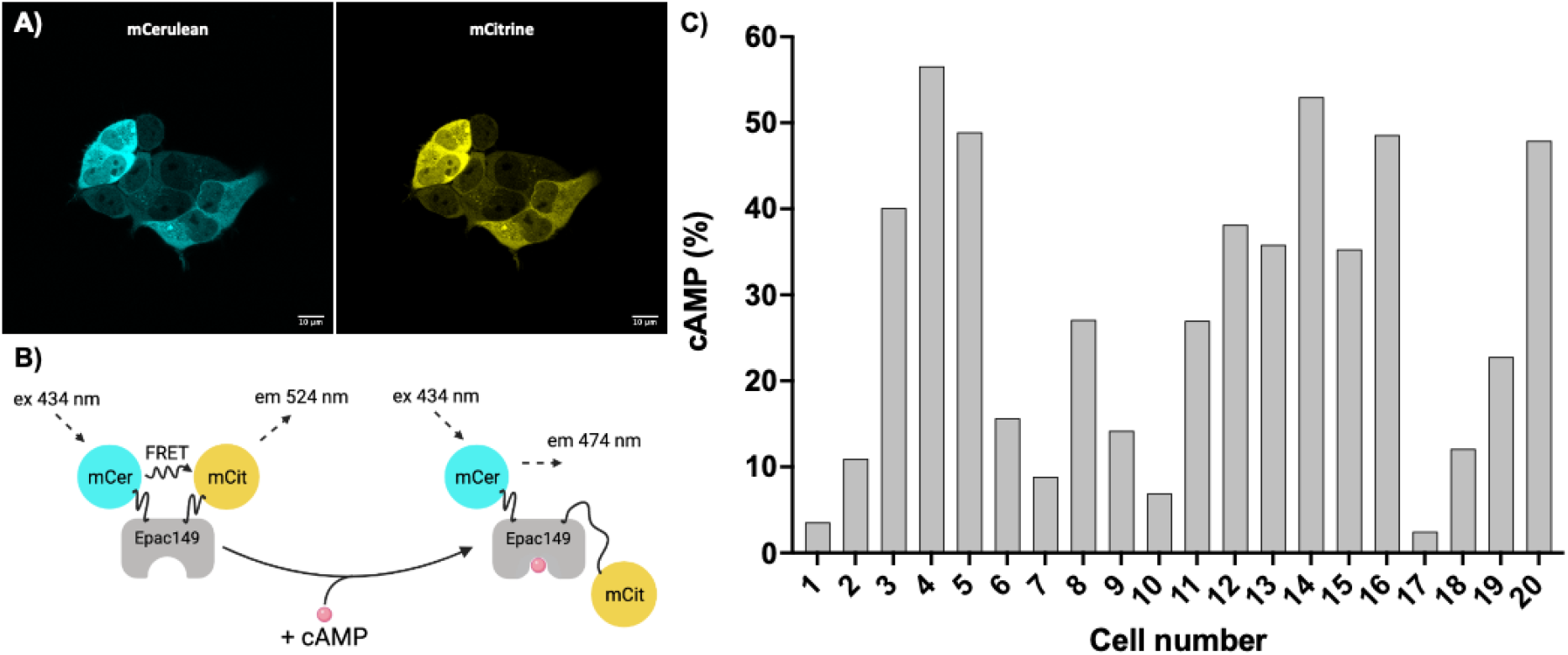
Detection of cAMP response in HEK293 cells expressing the Epac149 biosensor. **A)** Representative LSCM images of HEK293 cells expressing the Epac149 biosensor. The donor fluorophore mCer is shown in cyan and the acceptor fluorophore mCit is shown in yellow. **B)** Schematic representation of the Epac149 biosensor. The sensor is based on a scaffold of the Epac1 that is flanked by two fluorescent proteins, mCer and mCit. **C)** Increase in intracellular cAMP levels of twenty living HEK293 cells expressing the Epac149 biosensor. The cells were stimulated with 100 µM forskolin to induce maximum cAMP production. While the intracellular cAMP levels in some cells increased up to 55 %, the intracellular levels of cAMP barely increased in some cells.

### Quantitative interpretation of intracellular cAMP levels in single cell confocal microscopy images requires a minimum level of Epac149 biosensor expression

To examine why forskolin in some cases induced a high response and in other cases only induced a minor cAMP response, the responses of 54 cells, imaged in the same conditions as above, were plotted as a function of the biosensor expression level in each cell, shown in **Figure 2A**. We observed a dependence of the increase in intracellular cAMP levels on the biosensor expression level. In the case of cells expressing low amounts of biosensor, we observed a linear increase of the cell response with the amount of expressed biosensor, whereas cells expressing higher amounts of biosensor all showed similar responses after reaching a plateau. This suggests that a minimum amount of biosensor is needed to detect the full cellular response, i.e. to benefit from the full dynamic range of the biosensor chosen.

**Figure 2.**
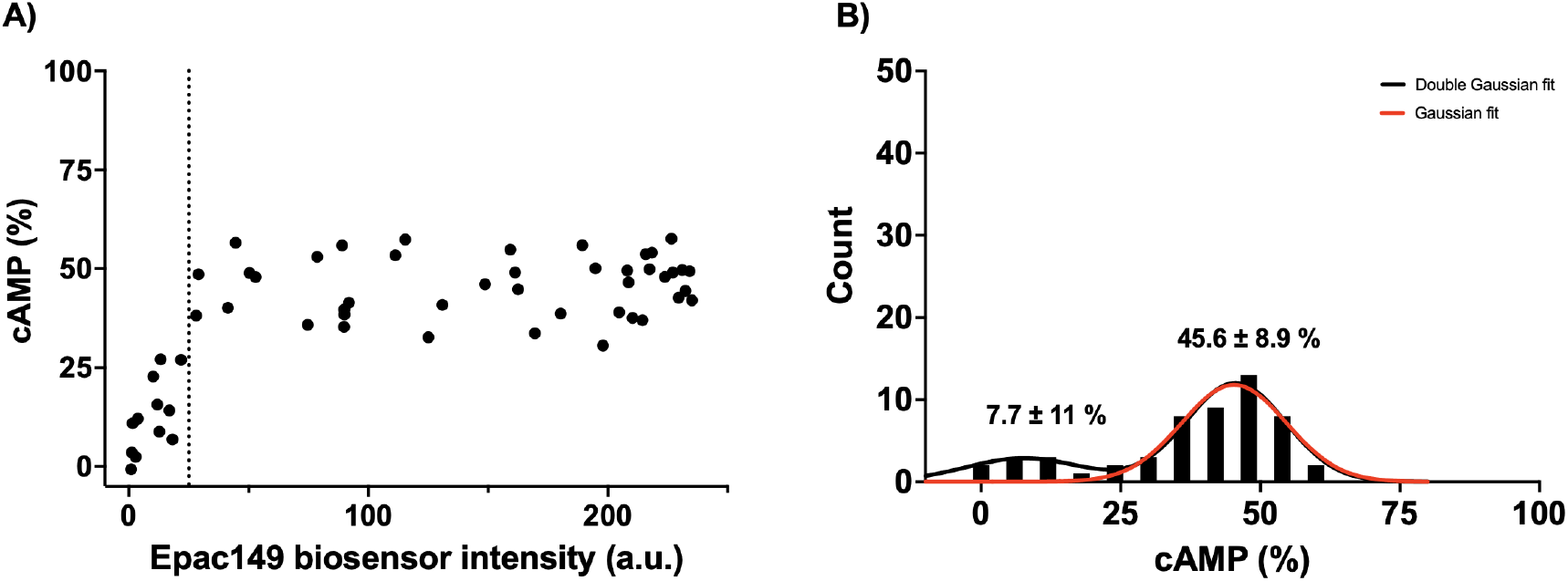
Maximum cAMP response of HEK293 cells transiently transfected with the Epac149 biosensor. **A)** Maximum cAMP response of transiently transfected HEK293 cells after stimulation with 100 µM forskolin. At low expression levels of the biosensor, the cAMP response of the cells increases linearly with an increase in biosensor expression level. As the expression level of the biosensor increases, the cAMP response of the cells stops increasing and hits a plateau. The mean increase in cAMP levels of all cells was calculated to be 38.3 ± 16 %. Each data point corresponds to a single cell, n = 54. **B)** Distribution of the data points from A). The data points are distributed in two distinct populations. One population of cells with a poor cAMP response, and one population of cells with a proper cAMP response. The distribution was fitted to a single Gaussian function and a double Gaussian function. The single Gaussian fit revealed a mean value of the cAMP response of 45.3 ± 9.4 %. The double Gaussian fit, however, revealed two mean values, one for each subpopulation of cells, of 7.7 ± 11 % and 45.6 ± 8.9 %.

The analysis of the distribution of the cAMP level increases, shown in **Figure 2B**, supports the notion of more than one distinct subpopulation of cells. A double Gaussian function was used to describe the distribution of data points, enabling an accurate fit of the whole distribution. From a statistical point of view, that suggests the presence of two subpopulations of cells. In this case, one subpopulation of cells with insufficient expression level of biosensor and a poor response to forskolin, and another subpopulation of cells with a higher expression level of the biosensor and a proper response to forskolin. From the double Gaussian fit, it was possible to extract the mean increase in intracellular cAMP levels from the two subpopulations of cells. The mean value of the poorly responding subpopulation was 7.7 ± 11 %, however, the standard error of the mean larger than the mean value is not suitable for quantitative studies of cAMP levels. The second subpopulation showed a better response at 45.6 ± 8.9 %. The difference in the cellular response of the two subpopulations further suggests that cells with an insufficient expression level of the biosensor do not respond properly to stimulation with forskolin. However, when expressed in quantities larger than the maximum possible cAMP production of the cell the biosensor provides us with a tool for optimal detection of changes in intracellular cAMP levels in living cells. Selection of cells that express suitable amounts of biosensor is therefore crucial for a correct use of the biosensor and an accurate interpretation of the cellular response to a certain stimuli. Such a selection is possible when investigating the response of single cells, but not if the response is measured by ensembles of cells. Indeed, measurements based on the overall response of an ensemble of cells, either per microscope image or per well of microtiter plates, provide the average readout of the whole cell sample. Transiently transfecting the cells can result in a substantial variation in protein expression levels from sample to sample, and therefore also a substantial difference in the readouts. Consequently, interpretation of cell studies relying on transient transfection of the plasmid coding for the biosensor have to be very cautious as the quantitative analysis will depend on the quality of the transfection, i.e. on the amount expressed in each cell composing the sample (or population).

### Changes in intracellular cAMP levels are independent of Epac149 biosensor expression levels in stable HEK293-Epac149 cells

We created a stable cell line with constitutive expression of the Epac149 biosensor in mammalian HEK293 cells and performed equivalent experiments using this cell line instead of transfecting the Epac149 biosensor into cells to circumvent the insufficient expression levels of the biosensor in some cells. In a similar manner as the previously described experiment, the HEK293-Epac149 cells were imaged in absence of ligand. After imaging, the cells were stimulated with 100 µM forskolin and imaged again 15 minutes after the addition of forskolin, shown in **Figure 3A**. This experiment showed that even in a stable monoclonal cell line the expression level of biosensor varies by more than 8-fold between different cells. However, all expression levels found in this cell line were higher than the threshold for insufficient expression levels we observed in transiently transfected cells. While the increase in intracellular cAMP levels depended on the biosensor expression in the transiently transfected cells, this was not the case for the stably expressing cells.

**Figure 3.**
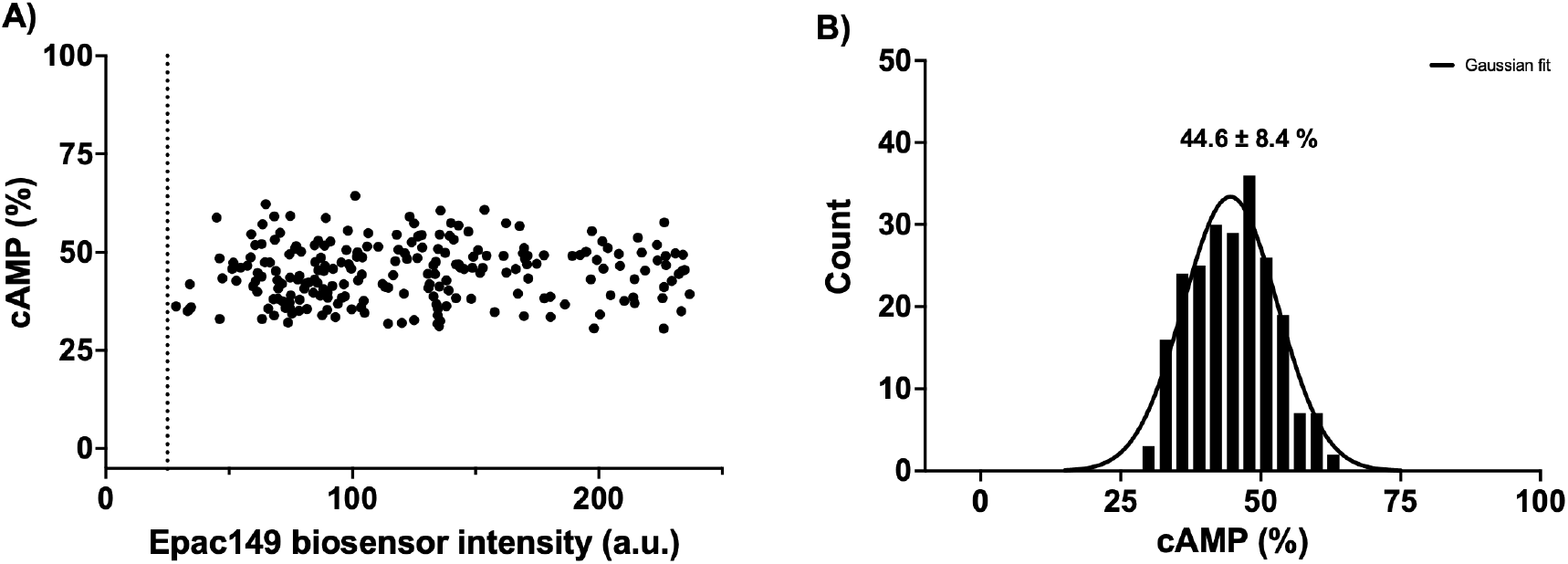
Maximum cAMP response of HEK293-Epac149 stable cell line. **A)** The maximum cAMP response of the HEK293-Epac149 stable cell line after stimulation with 100 µM forskolin. The response of the cells is independent of the expression level of the Epac149 biosensor. Each data point corresponds to a single cell, n = 223. **B)** Distribution of data points from A). The data points are distributed as a single distinct population. The distribution of cAMP responses was fitted with a single Gaussian function from which the mean cAMP response of the stable HEK293-Epac149 cell line was extracted to be 44.6 ± 8.4 %.

We binned the data and plotted the distribution of the cAMP levels increase, as shown in **Figure 3B** and found only a single population of cells. Therefore, we fitted the distribution to a Gaussian function and extracted the mean increase in cAMP levels as 44.6 ± 8.4 %. This was equivalent to the mean increase in cAMP levels of the properly responding subpopulation of the transiently transfected cells. As a result, it means that all the cells in the stable HEK293-Epac149 cell line express quantities of the biosensor in the range where we can make quantitative studies since the sensor responds linearly in all cells.

### Stability of the Epac149 biosensor in stable HEK293-Epac149 cell line

Time-resolved measurements of intracellular cAMP levels are an important aspect of obtaining novel information about cell signaling. For such experiments, it is required that the biosensor remains stable for a long time during acquisition. Therefore, we tested the stability of the biosensor in the HEK293-Epac149 stable cell line by conducting measurements over time. The cells were imaged every 2 minutes for 10 minutes prior to addition of forskolin followed by imaging every 2 minutes for 50 minutes, as shown in **Figure 4**. During the initial 10 minutes before addition of forskolin, the cAMP response of the cells remained stable around 0 %. After addition of forskolin, the cAMP response of the cells rapidly increased to 37.9 ± 2.9 % where it remained stable for at least 40 minutes.

**Figure 4:**
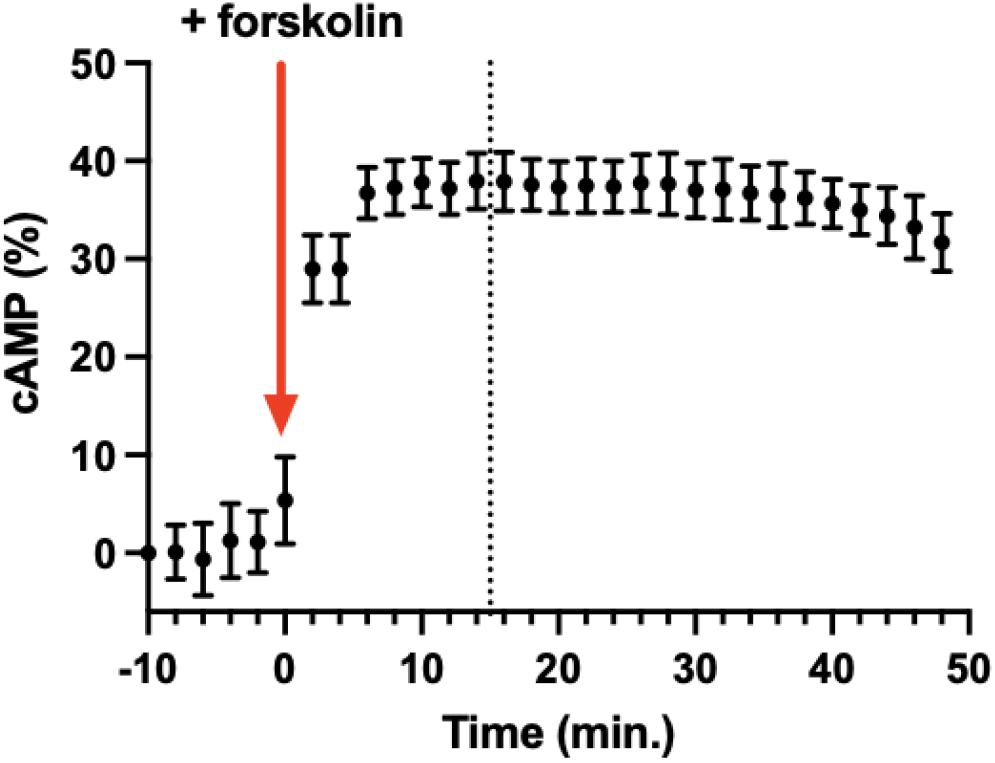
Time-resolved cAMP response of HEK293-Epac149 stable cell line. Maximum cAMP response of HEK293-Epac149 stable cell line over time. The cells were imaged every 2 minutes for 10 minutes prior to addition of ligand. Forskolin was added at a concentration of 100 µM (red arrow) and the cells were imaged every 2 minutes for 50 minutes. Prior to addition of forskolin, the cAMP response of the cells remained stable around 0 %. After addition of forskolin, the cAMP response of the cells rapidly increases to 37.9 ± 2.9 % and remains stable for 40 minutes. Each data point is the mean value of 4 cells measured over time ± SD. The dotted line at 15 minutes indicates the time point of measurements conducted in Figure 1, 2, and 3.

### Dose response curves established based on single-cell cAMP measurements by quantitative fluorescence microscopy

We evaluated if the poorly responding cells, eg. cells with insufficient expression levels of the biosensor, have an impact on the dose response curves when GPCRs are stimulated with a ligand. First, we either transiently transfected the HEK293-Epac149 stable cell line with SNAP-β_1_AR to express the β_1_-adrenergic receptor at the cell surface or transiently transfected the iSNAP-β_1_AR stable cell line with the biosensor. The receptor contained a SNAP-tag, which allowed us to chemically couple a small fluorophore on the extracellular N-terminus. The cells were imaged prior to addition of the β_1_AR partial agonist denopamine to evaluate the intracellular cAMP levels in the absence of ligand. After imaging, the cells were stimulated with varying concentrations of denopamine followed by imaging 15 minutes after addition of denopamine, as shown in **Figure 5D**. The cAMP response of iSNAP-β_1_AR cells transiently transfected with the biosensor, shown in **Figure 5D, gray** showed a 54.7 ± % response compared to the maximum induced by forskolin with a LogEC_50_ value of −9.20 ± 0.31. In comparison, when we selected only cells with sufficient biosensor expression levels we observed an increase in the cAMP response induced by denopamine to 79.7 ± 5.0 % of the maximum induced by forskolin with a LogEC_50_ value of −8.75 ± 0.18, shown in **Figure 5D, red**. Additionally, we measured the cAMP response of the stable HEK293-Epac149 cell line transiently transfected with SNAP-β_1_AR in which all cells express sufficient quantities of the biosensor, shown in **Figure 5D, blue**. Here, we found a cAMP response induced by denopamine of 81.2 ± 4.4 % of the maximum response induced by forskolin with a LogEC_50_ value of −8.57 ± 0.24. This response was similar to the response of the selected iSNAP-β_1_AR cells.

**Figure 5.**
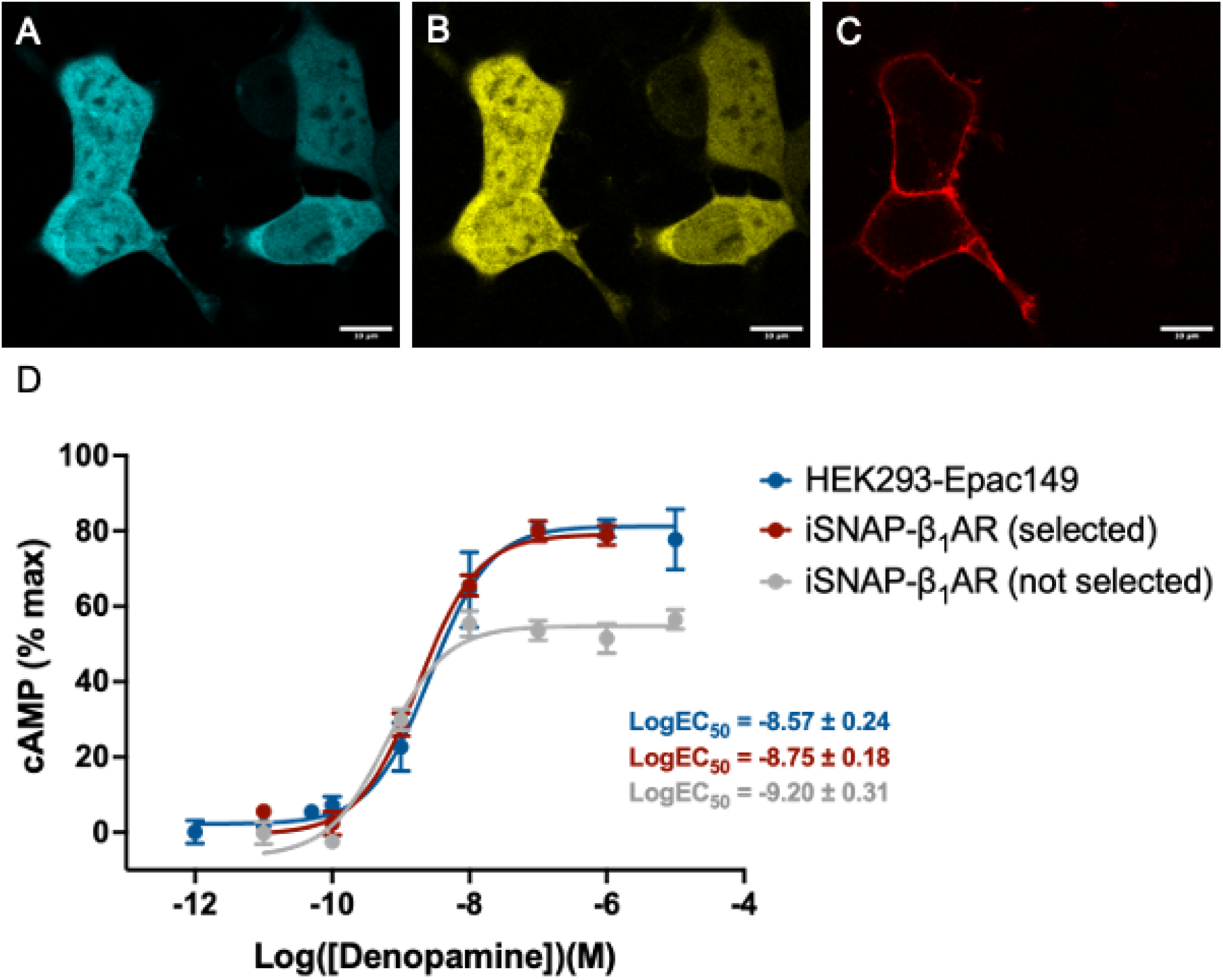
Dose response curves of HEK293 cells expressing β_1_ adrenergic receptor and Epac149 biosensor. Representative LSCM images of HEK293 cells expressing the Epac149 biosensor transiently transfected with SNAP-β_1_AR. **A)** Emission of mCer. **B)** Emission of mCit. **C)** Emission of SNAP-Surface AF647 labeled SNAP-β_1_AR. **D)** Dose response curves of HEK293 cells stimulated with the β_1_AR partial agonist denopamine. The cAMP response of iSNAP-β_1_AR cells transiently transfected with the Epac149 biosensor (gray) showed a cAMP response of 54.7 ± 3.4 % of the maximum response induced by forskolin. The LogEC_50_ value was extracted to be −9.20 ± 0.31. Including iSNAP-β_1_AR cells selected based on Epac149 biosensor expression levels only (red) increased the cAMP response induced by denopamine to 79.7 ± 5.0 % of the maximum induced by forskolin. The LogEC_50_ value was extracted to be −8.75 ± 0.18. The cAMP response induced by denopamine in HEK293-Epac149 stable cells was measured to be 81.2 ± 4.4 % of the maximum response induced by forskolin with a LogEC_50_ value of −8.57 ± 0.24. The data points of each curve represent the mean value of all individually measured cells ± SEM, n ≥ 3.

The consequence of insufficient biosensor expression in the measured cells is directly reflected in the dose response curves as both a lowered response and as a shift in the EC_50_ value. The iSNAP-β_1_AR cells that expressed biosensor in quantities large enough to properly detect the induced cAMP response showed a 1.46-fold increase compared to the cAMP response when not applying a post-selection of the cells. The stable HEK293-Epac149 cells showed a 1.48-fold increase in the response. The shift in LogEC_50_ value was 2.82-fold and 4.27-fold, respectively.

Characterization of biosensors and their performance in terms of dynamic range, sensitivity and linearity of the response is typically performed in solution using isolated sensors. In this case, the sensor is in excess compared to cAMP which ensures a linear response of the signal. However, confinement of the biosensor inside a cell can result in a deficit of the biosensor compared to the maximum possible cAMP production of the cell. In such a case the linearity of the measured signal is compromised which limits the quantitative interpretation of the measurement. To ensure correct working conditions, we propose two solutions. First, if the cells are transiently transfected with the biosensor, it is important to make sure that all cells express quantities of the biosensor suitable for optimal analysis of the cellular response in living cells. This solution is only relevant in the case of single cell measurements in which a post-selection can be applied in the analysis of the cells. However, if the assay is based on the use of microtiter plate readers, it is important to work with a well-characterized stable cell line. It is especially important in such an assay that all the cells express enough biosensor as post-selection inherently is not possible. Nonetheless, by applying the method we have shown here, microtiter plate reader assays can easily be adapted for measurements of cellular response in living cells.

## Conclusion

The assay we present here allowed detection of G_s_-coupled receptor activation by quantitative fluorescence microscopy in single living cells. When we investigated the cellular response to direct activation of the adenylyl cyclase by forskolin we observed an increase of the measured intracellular cAMP levels, that depended on the expression level of the Epac149 biosensor we used in cells transiently transfected with the biosensor. At low expression levels the linearity of the signal is lost due to a deficit of the biosensor compared to the maximum possible production of cAMP by the cell. At higher expression levels, however, the linearity was maintained as the biosensor is expressed in an excess. We created a stable cell line with constitutive expression of the biosensor to circumvent the insufficient expression levels of the biosensor in some cells. All cells in this stable cell line were observed to respond linearly to stimulation with forskolin.

We investigated the dose dependent cAMP response of cells expressing β_1_-adrenergic receptors by stimulation with the β_1_AR partial agonist denopamine. The consequence of insufficient biosensor expression was directly reflected in the dose response curves. Selection of transiently transfected cells with sufficient biosensor expression showed a 1.46-fold increase in the cellular response compared to the scenario without post-selection of cells. The stable HEK293-Epac149 cell line showed a 1.48-fold increase. Additionally, the LogEC_50_ values were shifted by 2.82-fold and 4.27-fold, respectively.

The approach we show here is applicable to any genetically encoded biosensor. It reveals the importance of careful characterization and control of the biosensor expression when investigating cellular responses. A small investment in the establishment of stable cell lines maintaining linearity of the biosensor signal allows the use of genetically encoded biosensor to be easily adapted for high-throughput screenings

## Notes

### Competing Interest Statement

The authors have declared no competing interest.

